# K978C CFTR restores essential epithelial function with greater efficiency than wildtype CFTR when expressed in CF airway cells

**DOI:** 10.1101/2022.08.10.503368

**Authors:** Maximillian Woodall, Robert Tarran, Rhianna Lee, Hafssa Anfishi, Stella Prins, John Counsell, Paola Vergani, Stephen Hart, Deborah Baines

## Abstract

Class Ia/b CFTR variants cause severe cystic fibrosis (CF) lung disease in ~10% of CF patients and are untreatable with small molecule pharmaceuticals. Genetic replacement strategies offer a potential cure for all patients but so far, have displayed limited efficiency *in vivo*.

We hypothesised that increasing protein abundance and/or activity of introduced CFTRs would more effectively restore function to CF bronchial epithelial cells (CFBE) in the presence of CF sputum (CFS) than wildtype (WT)-CFTR. We investigated codon optimised CFTR (hCAI), increased open probability CFTR (K978C) and codon optimised plus K978C (h^K978C) as candidates for gene therapy. Transfection of HEK293T with hCAI and h^K978C produced ~10-fold more CFTR protein than WT or K978C CFTRs. hCAI and h^K978C also displayed ~4-fold greater anion transport than WT in a halide-sensitive YFP quenching assay. However, functionality of modified CFTR cDNAs expressed in CFBE were profoundly different. 10% transduction of CFBE with K978C, compared to 22% transduction with WT, restored Cl^-^ transport to similar levels as that recorded from non-CF cells. K978C increased ASL height and pH more effectively than WT-CFTR, while hCAI and h^K978C had limited impact. Further investigation indicated that codon optimised CFTRs mis-localised in CFBE and compromised vectoral Cl^-^ transport.

These data provide further evidence that codon optimised CFTR cDNAs may be unsuitable for gene therapy practices that employ high activity promoters. However, increased activity CFTR cDNAs such as K978C, that potentially mimic the effect of potentiators, may provide more potent recovery of function than WT-CFTR cDNA in CF airways.

**Significance Statement:** Cystic fibrosis (CF) disease is associated with genetic malfunction of the Cl^-^ channel CFTR, leading to dehydration and decreased pH in the fluid lining the airways. Replacement of CFTR by gene therapy/gene editing offers potential therapeutic benefit but efficiency is poor. We show that gain of activity K978C CFTR under the control of a high activity promoter fully restored Cl^-^ transport, hydration and pH to CF bronchial epithelial cells (CFBE) in the presence of CF sputum and more efficiently than wild type CFTR. Codon optimised forms of CFTR were much less effective and proteins were mis-localised/mis-processed in CFBE. Thus, K978C could offer improved therapeutic potential.

## Introduction

Precision medicine using pharmacological drugs has had great success in treating cystic fibrosis (CF) variants with recoverable cystic fibrosis transmembrane regulator (CFTR) protein. However, there are no effective treatments currently available for class Ia (no mRNA) and class Ib (no protein) CFTRs, responsible for ~10% of CF causing variants (1). Developing gene therapy and gene editing techniques offer potential treatment for all CF mutations. Gene editing is in its infancy with proof-of-concept studies moving forwards. Gene therapy clinical trials for CF have had limited success. One barrier is that the efficiency of techniques to replace/correct CFTR in the airway *in vivo* is notoriously low. It is generally accepted that 10-25% of cells expressing endogenous CFTR will restore sufficient anion transport to reduce disease progression (2–6). Research in vitro suggests that transduction of 6-10% CF cells with exogenous WT CFTR cDNA generated Cl transport properties similar to normal cultures (7). However, a direct functional comparison of % cells expressing endogenous CFTR vs exogenous CFTR in human bronchial epithelial cells while taking into account the possible interference of the CF luminal environment on epithelial function and the activity of CFTR has not been documented.

Repeat dosing to improve transduction efficiency has led to detrimental side effects associated with immune responses to gene therapy. Codon optimisation of CFTR has been used to reduce immunogenicity of the introduced DNA by depletion of cytosine-phosphorothioate-guanine (CpG) dinucleotides (8, 9) Codon optimisation can also increase translation of the subsequent mRNA into protein by replacement of rare codons which can slow/terminate translation with those that match the most prevalent tRNAs in the host cell (10, 11). The replacement of the rare codons in WT-CFTR increased protein production when expressed in Fisher Rat Thyroid (FRT) cells, but the function of the codon optimised WT-CFTR protein was not fully investigated (11).

Exploration of CFTR structure and function relationships has identified several gain-of-function mutations (12). One gain of activity mutation, K978C within cytosolic loop 3 of CFTR, has been suggested to operate in a similar mechanism to the therapeutic drug VX-770 (13–15).

There is limited research on the function of codon optimized or K978C CFTRs in CF bronchial epithelial cells (CFBE) grown at air-liquid-interface, the gold standard for preclinical airway model systems (16). Therefore, we first established how CFTR-dependent Cl-secretion (measured as short circuit current (I_sc_)) and ASL height changed with the amount of endogenously expressed WT-CFTR using a CFBE and non-CF bronchial epithelial cell (NHBE) co-culture model. We then tested the hypothesis that transduction with an exogenous codon optimised form of CFTR or K978C-CFTR or codon optimisation together with K978C-CFTR under the control of a high promoter would reduce the efficiency required to restore CFTR activity to levels reported to be sufficient to ameliorate disease progression. We conducted all measurements in CFBE exposed to CF sputum to better mimic the CF environment and because our previous work indicated that acute exposure to CF sputum decreased CFTR function (17).

## Results

### CFTR activity is associated with % NHBE in co-cultures in the presence of CF sputum

Male (XY) or female (XX) donor CFBE or NHBE were isolated from bronchial regions of CF and non-CF lungs, counted and mixed at 10%, 20%, 50% and 75% (NHBE:CFBE) as represented (Fig. 1A). Markers of epithelial differentiation including cilia (a-tubulin) and tight junctions (phalloidin) were present in all mixed-cultures (Fig. 1B). The final percentage of NHBE:CFBE in the co-cultures, as analysed by droplet digital PCR for AMEL-X or AMEL-Y, was different to the original seeding percentage. This appeared to be donor-specific and independent of the CF mutation or whether the donor was male or female (Fig. 1C).

**Figure 1.**
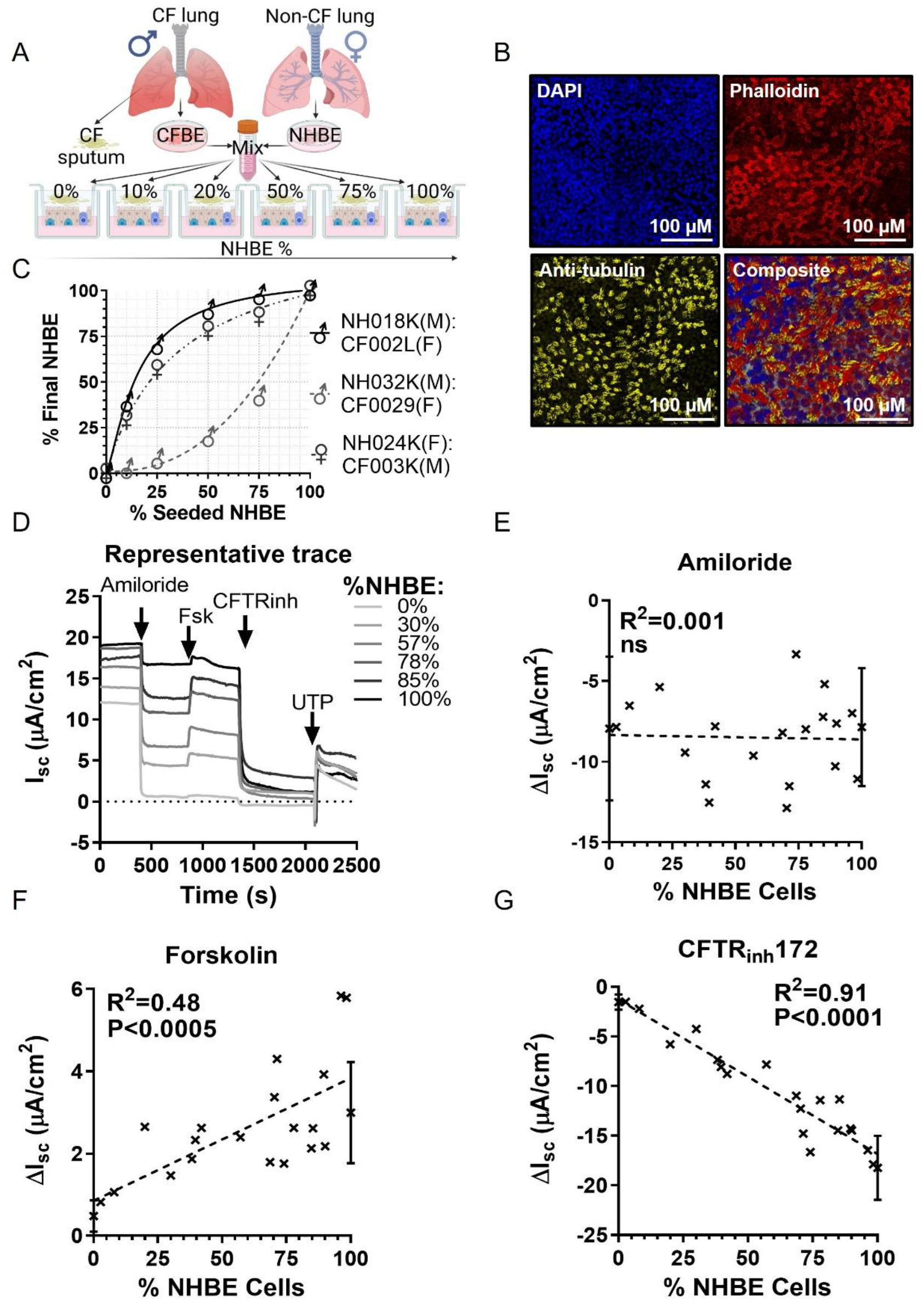
Characterisation of the NHBE:CFBE mixed culture model. **(A)** Schematic diagram of the NHBE:CFBE mixed culture model. CFBE and NHBE from 3 male (XY) or 3 female (XX) donors were combined in ratios of 10%, 25%, 50% and 75% NHBE:CFBE and seeded onto semipermeable membranes. (B) Confocal microscope images (XY view from compressed z-stack) of an example 50% NHBE:CFBE, stained with phalloidin (red; F-actin), Anti-β-tubulin (yellow; cilia), DAPI (blue; nuclei) and composite image. Scale bars are shown in the bottom right-hand corner. (C) Percentages of AMELX and AMELY regions of genomic DNA from three different mixes of male (M) and female (F) NHBE or CF donors (see SI, Table 1) post-functional analysis, as determined using droplet digital PCR. Data were normalised to values from 100% male or100% female cultures and 28 samples were run in duplicate. Each data point represents the mean ± SD (error bars are smaller than the height of symbols). (D) Representative I_sc_ traces from NHBE:CFBE after incubation with CFS and addition of specific activators and inhibitors of ion transport (as below). I_sc_ plotted against % NHBE:CFBE (n=26 and 6 individual donors) for (E) ΔI_sc_ after addition of amiloride (100 μM), (F) forskolin (10 μM) and (G) CFTR_inh_172 (10 μM). All drugs were added apically with exception of forskolin which was also added basolaterally. Data were subject to linear regression. The R^2^ values and P value for slope significantly different from zero, are shown on the graphs. ns., not significantly different.

We investigated bioelectric properties of NHBE:CFBE co-cultures in the presence of CF sputum (Fig. 1D). Responses to amiloride, forskolin, CFTR_inh_172 and UTP provided evidence that functional epithelial Na^+^ channels (ENaC), CFTR and calcium-activated chloride channels (CaCC) were present. We observed a linear correlation of change in short circuit current (ΔI_sc_) with % of NHBE in the co-culture in response to forskolin (R^2^=0.48; P<0.0005) (Fig. 1F) and CFTR_inh_172 (R^2^=0.91; P<0.0001) (Fig. 1G), but not with amiloride (Fig. 1E) or UTP (S1) (n=26, from 6 individual donors).

### Expression and function of CFTR cDNAs in HEK293T cells

All constructs (Fig. 2A) produced CFTR protein, visible as the mature, complex glycosylated ~170kDa protein (band C), and immature core glycosylated ~140kDa protein (band B) by western blot (Fig. 2B). The YFP protein product in co-transfected HEK293T cells was observed at ~22kDa. Codon optimised hCAI and h^K978C produced more CFTR protein than WT or K978C, with hCAI producing ~14-fold more than WT (p<0.05; n=3) (Fig. 2C). K978C-CFTR transfection caused a loss of adherence of HEK293T and so these cells could not be used for downstream analysis other than lysis for protein collection. A YFP quenching assay was employed to determine function of CFTRs (Fig. 2D-F). There was little decrease of YFP fluorescence in the presence of DMSO (vehicle) (Fig. 2D). Addition of CFTR activator forskolin to WT, hCAI and h^K978C transfected HEK293T cells produced a decrease in YFP fluorescence (Fig. 2E), with h^K978C displaying significantly greater maximal rate of I^-^ entry compared to hCAI (0.3 mM/s and 0.23 mM/s, respectively p<0.05; n=3), and WT-CFTR (0.06 mM/s, p<0.001, n=3) (Fig. 2G). Prior exposure of hCAI to potentiator VX770 increased quenching of YFP fluorescence (n=3, p<0.01) producing a similar maximal rate of I^-^ entry to h^K978C (Fig. 2F and G). WT quenching of YFP was also increased >2-fold (0.057±0.011 to 0.15±0.012, p<0.01; n=3) (S2) though this remained at a lower rate than hCAI or h^K978C (0.15 mM/s, 0.30 mM/s and 0.32 mM/s respectively, p<0.05; n=3) (Fig. 2G).

**Figure 2.**
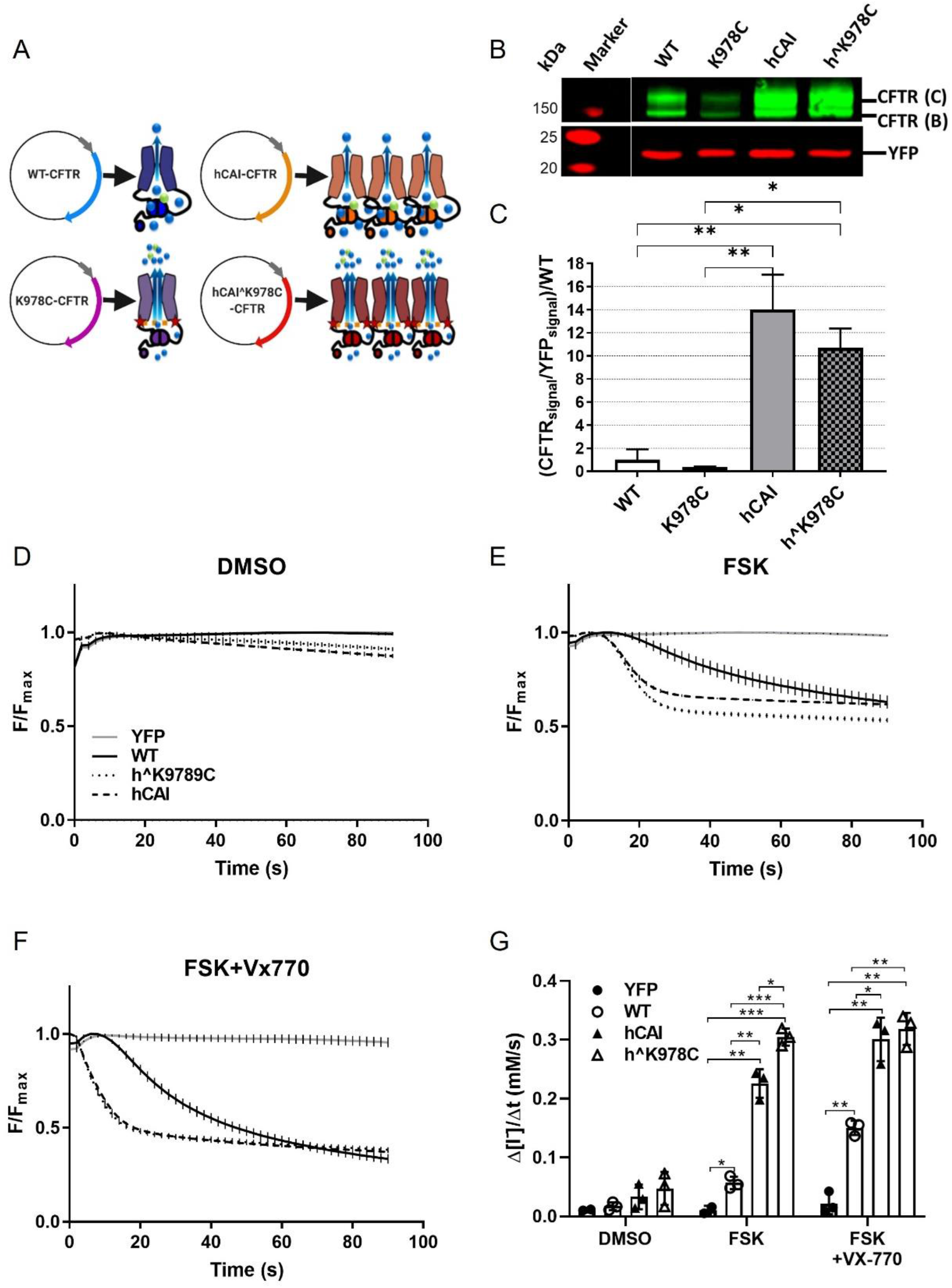
Changes to CFTR sequences affect protein production and function in transfected HEK293T cells. (A) Simplified plasmid maps for each version of CFTR cloned down-stream of a CMV promoter (grey): wild type in blue (WT), K978C mutation in purple (K978C), high codon adaption index CFTR in orange (hCAI) and hCAI with the K978C mutation in red (h^K978C) with accompanying cartoon representing abundance and activity of protein produced by each plasmid in HEK293T cells: Arrows indicate anion movement (blue and green spheres), K978C mutation (red stars). (B) Example of a western blot of protein extracted from HEK293T cells co-transfected with YFP and CFTR cDNAs as shown. CFTR bands C and B (green) and YFP (red). Molecular weight markers (also red) are shown to the left-hand side of the image. (C) Quantification of CFTR protein abundance is shown below the blot. Data are shown as mean (CFTR (band C + B) /YFP) normalised to WT to standardise between blots. Data are shown as means +/- SD. Treatments were compared by one-way ANOVA with Tukey’s post hoc analyses; Significantly different as shown *: p<0.05; **: p<0.01; n=3. (D) YFP fluorescence quenching over time shown for HEK293 cells transfected with YFP alone (YFP) or co-transfected with CFTR cDNAs (as described above) in the presence of DMSO (vehicle) (D), Forskolin (FSK) (E), FSK plus the CFTR potentiator VX770 (FSK+VX770) (F). The maximal rate of I^-^ entry (Δ[I^-^]/Δt) summarised for CFTR cDNA and conditions shown in (G). Data are shown mean +/- SD with individual data points. CFTR function was compared by two-way ANOVA with Tukey’s post hoc analyses; Significantly different as shown *: p<0.05; **: p<0.01; ***: p<0.001, n=3.

### Function of CFTR cDNAs in CFBE

CFTRs upstream of GFP separated by a self-cleaving T2A peptide were transduced into CF cells (W1282X/R1162X or ΔF508) using lentivirus in the presence of CF sputum. Transduced CFBE-maintained transgene expression for 3-4 weeks and terminally differentiated ciliated cells were present (Fig. 3A and B). I_sc_ was measured in the transduced CFBE cells (Fig. 3C). The summarised data from all donors and transductions showed that ΔI_sc_ in response to CFTR_inh_172 increased in a linear fashion with % transduction of K978C, WT and hCAI CFTRs (R^2^=0.95, 0.93 and 0.57, P<0001, n=8-12 from 3 donors respectively). A similar correlation was observed with basal and forskolin-sensitive I_sc_ (S3. C). h^K978C was poorly correlated with any of these functions. Transduction of K978C increased CFTR_inh_172 ΔI_sc_ >2-fold compared to WT and regression analysis indicated that 9.8% transduction of K978C and 20.4% of WT would be sufficient to restore CFTR mediated I_sc_ to NHBE levels of −17μA/cm^-2^ (Fig. 3D). Codon optimised CFTRs produced smaller CFTR_inh_172 ΔI_sc_ than WT (Fig. 3D). There was little or no correlation of % transduction of any CFTR cDNA with amiloride-sensitive I_sc_ or UTP stimulated I_sc_ (S3).

**Figure 3.**
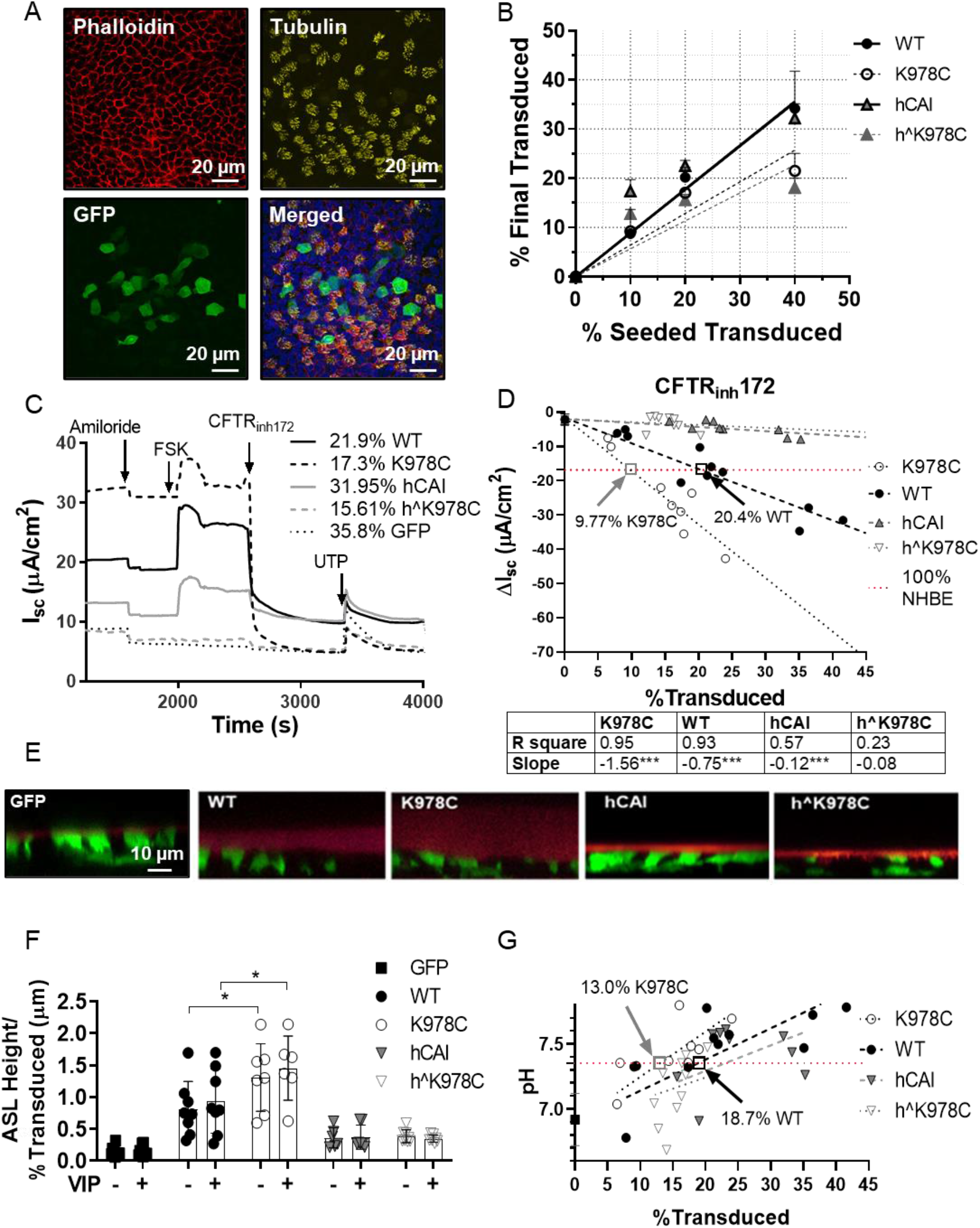
Characterisation of CFTR transduced CFBE. **(A)** Confocal microscope images of GFP CFTR transduced CFBE cultures, immunostained with phalloidin (F-actin; red), anti-β-tubulin (cilia; yellow), GFP (transduced cells; green) and a merged image which also includes nuclei stained with DAPI (blue). (B) Relationship between % transduced cells at seeding and % transduced after culture at air-liquid interface (ALI) for the different CFTR cDNAs. (C) Example I_sc_ traces from transduced CFBE cultures (% transduction for individual CFTR cDNAs are given) after incubation with CFS and addition of amiloride (10 μM), forskolin (10 μM), CFTR_inh_172 (10 μM) and UTP (10 μM). All drugs were added apically except forskolin which was bilateral. (D) ΔI_sc_ in response to CFTR_inh_172 plotted against % transduced for each CFTR cDNA. Dotted lines show linear regression with R^2^ values and significance values for variation of the slope from zero for each CFTR cDNA are shown in the table below *: p<0.05; **: p<0.01; ***: p<0.001. (n=8-12 from 3 CF donors). The red dotted horizontal line depicts value for ΔI_sc_ CFTR_inh_172 in 100% NHBE and labelled arrows show % transduction of WT and K978C required to reach this value. (E) Representative XZ images of airway surface liquid (ASL) labelled with dextran (red layer) overlying transduced CFBE cultures following incubation with CFS. Transduced cells express cytosolic GFP (green). (F) Summary of ASL height in μm/% transduced measured in CFBE transduced with CFTR cDNAs in the presence of CFS with (+) and without (-) stimulation with vasoactive intestinal peptide (VIP). Data are shown as mean +/- SD with individual data points for n=6-11 from 3 CF donors. Treatments were compared by two-way ANOVA with Tukey’s post hoc analyses; Significantly increase compared to WT *: p<0.05. (G) ASL pH plotted against % transduced CFBE for CFTR cDNAs. Dotted lines show linear regression for each CFTR cDNA (n=8-11 from 3 CF donors). *: p<0.05; **: p<0.01; ***: p<0.001. The solid horizontal line marks physiological pH (7.35) and vertical lines show the % transduction of WT and K978C required to reach this value.

Airway surface liquid (ASL) height per % transduced was greatest in CFBE transduced with K978C both without (1.31±0.53) and with stimulation with vasoactive intestinal peptide (VIP) (1.45±0.50) (Fig. 3E *and* F). The ASL height generated was significantly greater (~1.6 fold) than that produced by WT CFTR (-VIP: 0.81±0.43; +VIP:0.93±0.50) (p<0.05, n=6-8 from 3 CF donors respectively) (Fig. 3F). hCAI and h^K978C transduced cultures had no significant effect on ASL height compared to GFP alone (Fig.3F). Similar to our previous findings there was no significant difference in ASL height with or without stimulation with VIP in the presence of CFS.

ASL pH in CFBE transduced with GFP alone was 6.9±0.2. pH increased to physiological levels (7.35 pH) in CFBE with 13.0% transduction of K978C (n=8 from 3 donors) and 18.7% WT (n=11 from 3 donors) (Fig. 3G). hCAI and h^K978C transduction was less efficient at restoring ASL pH to physiological levels (n=9-10 from 3 donors) (Fig. 3G).

### Codon optimised CFTRs are miss-localised in CFBE

Transduction of CFBE with all CFTR cDNAs produced distinct CFTR proteins, mature (band C ~170kDa), immature (band B ~140kDa) (Fig. 4A). As expected hCAI produced the most CFTR protein per transduced cell (p<0.001, n=5-7 from 3 donors). However, both h^K978C produced less CFTR protein than WT (p<0.05, n=4-7 from 3 donors). The abundance of band C to B (ratio: 5.1-5.2) were significantly greater across all forms of CFTR (p<0.05, n=4-7 from 3 donors) other than h^K978C (Fig. 5B). The amount of bicistronic GFP in the cell broadly followed the same pattern except that GFP for codon optimized h^K978C was higher than K978C (p<0.05, n=4-7 from 3 donors) (S4). These data indicate that CFTRs containing K978C may be subject to increased degradation/turnover at the membrane.

**Figure 4.**
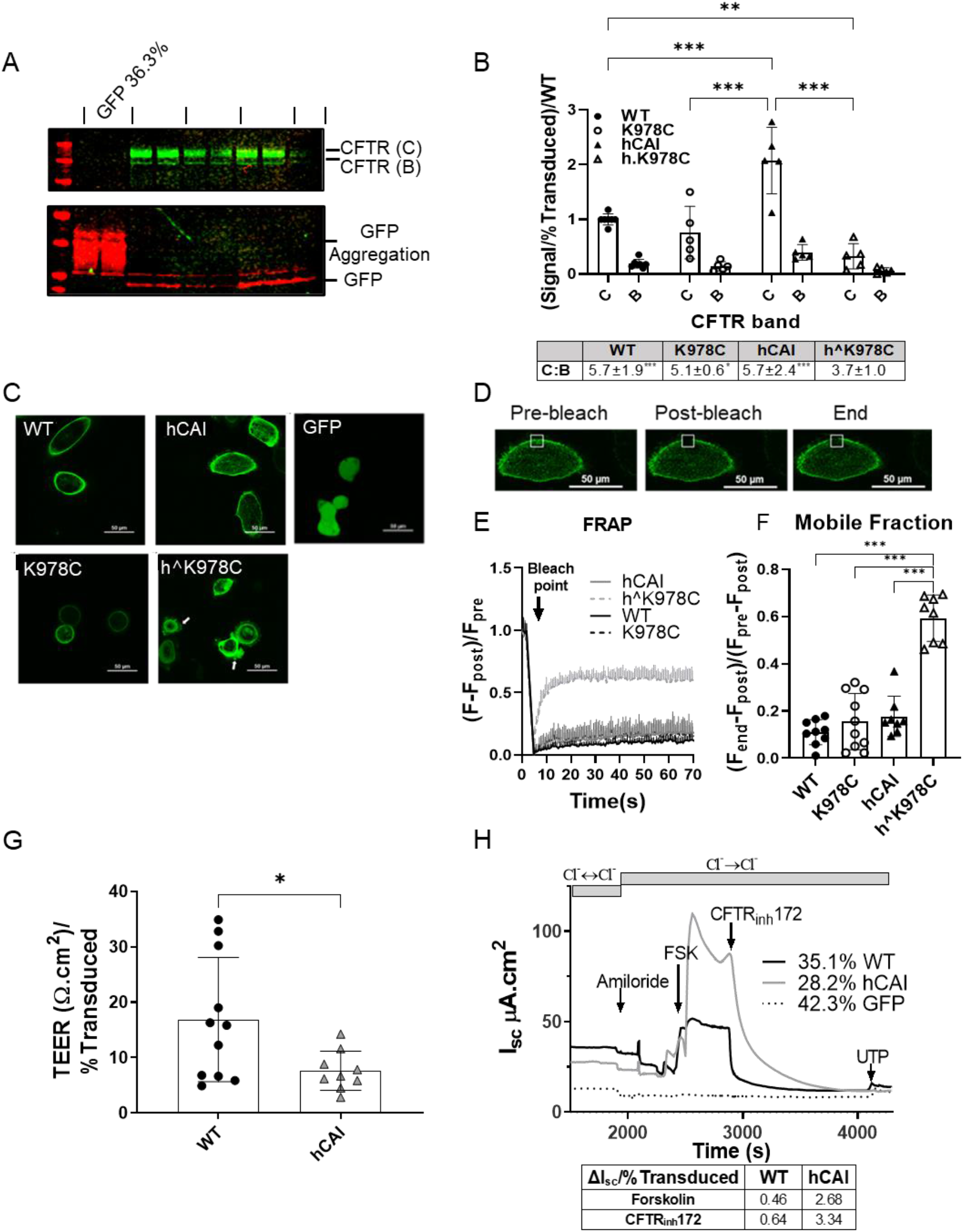
Overexpression of CFTR protein in CFBE cultures results in mis-localisation. (A) Example western blot of soluble protein fraction from CFBE cultures transduced with CFTR cDNAs or GFP alone (as labelled above wells). Blots were immunostained with anti-CFTR (CFTR 596) and anti-GFP. CFTR protein bands C and B (green), GFP (red) with molecular weight markers shown to the left of the blot (red). (B) Summary of analysis of relative fluorescence units (RFU) for CFTR bands C and B relative to % transduced and normalised to WT for each CFTR cDNA. Presented as mean ± SD with individual data points (n=4-7 from 3 donors). Treatments were compared to WT-CFTR by two-way ANOVA with Tukey’s post hoc analyses; Significantly different ** p<0.01, *** p<0.001. Ratio of RFU of bands B:C are shown below the graph. (C) Example of confocal microscope live images (x60, Ex:488/Em:507) of CFBE transduced with GFP linked to CFTR CDNAs or GFP alone. A 50 μm scale bar is shown at the bottom right of each image, white arrows show cell blebbing and (D) images taken during FRAP protocol, showing point of highest fluorescence in a cell (Pre-bleach), immediately after bleaching (Post-bleach) and 65 seconds after bleaching (End). The region of interest (ROI) is within the white square, the 50μm scale bar is shown at the bottom right of each image. (E) Kinetic plot displaying fluorescence (F) change over time for a ROI on the periphery of transduced cells. The time of bleaching with 100% 488 laser (Bleach point) is marked with an arrow. (F) Summary of the mobile fraction (Mf) ratio for different GFP linked CFTRs. Data are shown as mean +/- SD with individual data points. Treatments were compared by one-way ANOVA with Tukey’s post hoc analyses. Significantly different as shown; **: p<0.01; ***: p<0.001. (n=8-13 from 1 CF donor: DF508/DF508). (G) TEER per %Transduced for WT and hCAI CFTR cDNAs. Treatments were compared by unpaired t test; Significantly different * p<0.05 (n=8-12 from 3 CF donors). (H) I_sc_ traces from CFBE transduced with K978C, WT or hCAI or GFP alone before and after addition of a basolateral to apical Cl^-^ gradient and specific activators and inhibitors of ion transport, amiloride (10 μM), forskolin (10 μM), CFTR_inh_172 (10 μM), and UTP (10 μM). All drugs were added apically with exception of forskolin which was added bilaterally. Direction of chloride concentration gradients (using physiological Cl^-^ and Cl^-^ free buffers) are shown and the duration of application is presented as grey bars. ΔI_sc_/% transduced for ΔI_sc_ in response to forskolin and CFTR_inh_172 are shown in a table below the graph.

GFP fusion to the N-terminus of CFTR was used to further investigate localisation of CFTRs in CFBE BMI-1. CFBE BMI-1 were used for this assay due to their extended proliferative capacity, allowing for reproducible acquisition of fluorescence recovery after photobleaching (FRAP) data across cell passages. WT and K978C GFP fluorescence were clearly localised to the cell membrane, hCAI showed increased membrane and intracellular fluorescence (S4. *C*). h^K978C was identified throughout the cell whilst cell blebbing, a characteristic marker of cell apoptosis was also observed (white arrows: Fig. 4C). FRAP of the highest fluorescent region of the transduced cells indicated that h^K978C displayed increased free, rapid diffusion compared to WT, K978C and hCAI which is defining of a non-membrane bound protein (p<0.001, n=8-10) (Fig. *5E* and *F)*. Thus, it is likely that the lack of h^K978C function was attributable to improper protein processing, membrane cycling and consequent effects on cell viability.

The lack of function associated with hCAI did not correlate with this data supporting increased abundance of a membrane bound protein. However, there is evidence that overexpression of CFTRs can result in basolateral expression of CFTR, reducing TEER and impeding vectoral ion transport in physiological conditions (29). In line with this theory, transduction with hCAI did cause a significant decrease in TEER compared to WT (p<0.05, n=8-12 from 3 CF donors) (Fig. 5G). Furthermore, we implemented an artificial basolateral to apical Cl^-^ gradient in the Ussing chamber to assess if hCAI transduced CFBE were able to facilitate Cl^-^ movement. Application of the chloride gradient did not immediately change I_sc_. However, unlike the response in symmetrical Cl^-^ solutions (Fig. 3C), in the presence of the Cl^-^ gradient, addition of forskolin produced a ΔI_sc_ surpassing that of WT (~5.8 fold) and this current was CFTR dependent as it was fully inhibited by CFTR_inh_172 (Fig. 4H).

## Discussion

The aim of this research was to carry out a direct functional comparison of % cells expressing endogenous CFTR vs exogenous CFTR cDNAs in human bronchial epithelial cells in the presence of CFS. There was a linear relationship between CFTR_inh_172-sensitive I_sc_ and % NHBE after exposure to CFS. These data contrast to findings where CFTR mediated Cl^-^ secretion plateaued with ≤50-75% CFTR expression in other mixed culture methods, reportedly limited by transporter driven basolateral Cl^-^ entry (3, 6, 29). We previously showed that the acute presence of CFS lowered CFTR-dependent I_sc_ in NHBE cultures (CFTR_inh_172 ΔI_sc_ was −18μA/cm^-2^ whereas others recorded 20-50μA/cm^-2^) (17). Thus, in our system CFTR_inh_172-sensitive I_sc_ may not be subject to Cl^-^ entry limitation. Furthermore, we aligned CFTR dependent function with final %NHBE measured using ddPCR amplification of sex specific chromosomal regions, accurate for quantification of population chimerism to 0.01% (20, 30). Previous studies used proliferation assays of primary cells prior to mixing or qPCR using primers to the ΔF508 region to estimate the final % of NHBEs in their models (3, 6, 29), a less sensitive method than ddPCR particularly as we showed cell proliferation varied widely, independent of sex or CF status.

As previously reported, in HEK293T cells, CFTR protein abundance from codon optimised cDNAs was greatly increased compared to WT (11). The replacement of lysine to cystine in both K978C and h^K978C decreased protein expression, similar to that reported for G551D/K978C compared to G551D expression in HEK293T cells (13). The halide transport function of the cDNAs corelated with protein abundance and the increased activity mutation. K978C is hypothesised to destabilise the inactive state of CFTR and lead to a 2-fold increase in open probability (Po) of the channel (13, 31). Interestingly, pre-treatment of hCAI with the CFTR activator VX-770 increased activity to the same level as h^K978C. VX770 is predicted to bind at the interface between the two transmembrane domains and induce a conformational change to stabilise the open, pre-hydrolytic state of CFTR (15, 32) increasing Po of WT-CFTR by ~2 fold (33, 34), a phenomenon that we replicated in our YFP quenching assay. Thus, our results support that K978C produces a similar outcome to that of CFTR potentiation by VX770, a mechanism well tolerated in people with CF.

In contrast, in CFBE, h^K978C proved the least effective functionally. However, K978C produced the largest CFTR_inh_172 sensitive I_sc_ and increased ASL height implying that K978C better facilitated Cl^-^ secretion than the other cDNAs. When compared to the NHBE:CFBE co-culture model only 10% transduction of K987C was required to restore CFTR function to that of a NHBE monolayer, half of that required for WT and consistent with the described increase in Po. ASL pH was also more effectively restored by K978C than WT (at 13% and 19% transduction respectively) indicating that the construct better promoted HCO3^-^ secretion. It was proposed that between 6-25% transduced-CFTR expression is generally accepted to produce normal function *in vitro* (4, 7) but approximately 10% of fully functional CFTR transcripts are necessary for prevention of lung disease progression *in vivo* (2). While we accept that endogenous and exogenous expression of CFTR is different, our data indicate it may be possible to achieve sufficient CFTR activity with less than 5% cellular expression of K978C.

Although codon optimisation of CFTR generated more protein and function in HEK293T cells, it did not translate to more effective vectorial Cl^-^ transport in CFBE. Our evidence indicated that protein expression and localisation of codon optimised CFTR in CFBE was disrupted. FRAP can be used to determine the rates of local protein turnover, identify mobile fractions, and demonstrate exchange between cellular compartments or lack thereof in live cells (35). In this assay, h^K978C displayed free, rapid diffusion, providing evidence that the protein was not membrane bound. While hCAI did exhibit evidence of lateral diffusion in the membrane-associated state, TEER was reduced and application of an artificial Cl^-^ gradient in Ussing experiments showed that hCAI was functional but mis-localised, impacting the vectorial transport of Cl^-^. In support of our findings, overexpression of CFTR has been shown to perturb polarisation of epithelial cells, localisation of membrane proteins and membrane potential which would all compromise CFTR mediated Cl^-^ transport (36–38). CFTR under control of a CMV promoter was observed to generate basolateral CFTR, while use of a low-activity K18 promoter resulted in greater Cl^-^ currents in differentiated CF airway epithelia (29). Thus, further work is necessary to determine if hCAI and h^K978C under the control of low activity promoters would be more effective at restoring essential epithelial characteristics to CFBE.

While ENaC has been described as a regulator of ASL height (24), we saw little change in amiloride-sensitive I_sc_ in our models. In our experiments, cells were passaged at least twice which may have impacted ENaC function (39). Nevertheless, the lack of altered ENaC activity in these cultures further supports the hypothesis that the large changes in ASL observed with K978C were predominantly mediated by CFTR function as opposed to changes in ENaC activity.

ASL pH was also more effectively restored by K978C than WT or hCAI. These data indicate that the constructs effectively promoted HCO3^-^ secretion. In differentiated CFTR knockout porcine models, transduction of >10% WT CFTR increased pH to physiological levels (6). Our data indicate that 13% and 19% transduction of K978C and WT respectively restored ASL pH in CFBE. Interestingly, hCAI and h^K978C transduction also restored ASL pH but less efficiently (>20%). One possible explanation is that the constructs increased HCO3^-^ secretion via other channels that interact with CFTR. For example, HCO3^-^ secretion is facilitated by SLC26 transporters that are expressed in the apical membrane together with CFTR (40).

In conclusion, we provide proof of principle that K978C CFTR was more effective at restoring essential epithelial characteristics to CFBE than WT CFTR at a lower transduction rate. Increasing CFTR protein using codon optimisation did not translate to better function in CFBE. Expression and function of CFTR in heterologous cell line systems, specifically HEK293T cells, while useful, does not replicate function in differentiated airway epithelial cells, and may be misleading in development of CF gene therapies.

## Materials and Methods

Further details are provided as Supplementary information.

### Primary Human Bronchial Epithelial Cell Culture

CFBE and NHBE were obtained with ethical approval obtained by the University of North Carolina at Chapel Hill Biomedical Review Board (protocol #03 1396). Cells were cultured on permeable supports and maintained at air-liquid interface (ALI) in a modified bronchial epithelial growth medium (18). Co-cultures were generated by mixing NHBE and CFBE of alternate sex (XX or XY) at 10%, 25%, 50% and 75% (non-CF:CF). Cultures were incubated with 20μl apically applied CF sputum 4 hrs prior to functional experimentation as described (17). Donor demographics are provided in *SI* Table 1. BMI-1 transduced Cystic fibrosis (DF508/DF508) bronchial epithelial cells (CFBE BMI-1) were generously supplied by Professor Stephen Hart (UCL, Institute of Child Health) (19).

### Droplet Digital PCR

ddPCR using primers for amelogenin-X isoform (AMELX) and amelogenin-Y isoform (AMELY) and probes with FAM and VIC reporters, respectively (AB-Bioscience, UK), were performed as previously described (20). Primer sequences are provided in *S*I, Table 7. FAM and VIC positive (+ve) droplets from each well of the PCR plate were measured and analysed using associated software QuantaSoft. These data are presented normalised with exclusively male cultures (n=10) set at 0% and female (n=10) set at 100%.

### Sputum preparation

Airway sputum samples were obtained as described in University of North Carolina (UNC) protocol (21). Donor demographics of induced sputum donors are provided in *S*I, Table 2 and are reported in (17). Preparation of samples are described in detail (17, 22).

### Immunohistochemistry

ALI cultures were fixed in 4% PFA, permeabilized and blocked (*SI*, Table 6) prior to overnight incubation at 4°C with primary antisera (α-tubulin, GFP) followed by visualization with Alexa Fluor 568 or Alexa Fluor 488. Membranes were counterstained with 1:50 phalloidin and 1:200 DAPI for 30 minutes at room temperature. Images were captured using a Leica SP8 confocal microscope with LAS AF (Leica) acquisition software (antisera and fluorophores: *S*I, Tables 4 and 5).

### Electrophysiological measurement

Transepithelial ion transport was measured using the Ussing chamber technique using symmetrical buffers (*S*I, Table 6) and the following drugs: amiloride, 100 μM (apical); forskolin, 10 μM (bilaterally), CFTR_inh_172, 10 μM (apical) and UTP 100 μM (apical) as previously described (17). For Cl^-^ gradient studies the apical chamber was replaced with Cl^-^ free Ussing Buffer (*S*I, Table 6). Data were analysed using Acquire and Analysis (version 1.2) software (Physiologic Instruments). All drugs/chemicals were obtained from Sigma-Aldrich.

### Airway surface liquid (ASL) height measurements

ASL was labelled with PBS (20 μl) containing 0.5 mg/ml of 10 kDa dextran-tetramethylrhodamine (Life Technologies, USA). Images were obtained before and 60 minutes after addition of basolateral VIP (100nM) to induce CFTR mediated secretion by using a Leica SP8 confocal microscope with a ×63/1.3 numerical aperture (NA) glycerol immersion lens in XZ-scanning mode as described (17, 24).

### Site-directed mutagenesis

CFTR with high codon adaptation index 39 (CAI) values (from www.jcat.de) and named hCAI was generously supplied by David Mueller (Department of Biochemistry and Molecular Biology, Rosalind Franklin University) (11). Primers containing the nucleotide sequence to alter WT CFTR with the K978C mutation (K978C), hCAI with the K978C mutation (h^K978C) or for T2A mutagenesis were designed using SnapGene software (*S*I, Table 7) and synthesised by Sigma Aldrich.

Site directed mutagenesis was carried out on an Eppendorf^®^ Mastercycler^®^ Pro Thermal Cycler using PhusionTM Hot Start II High-Fidelity PCR Master Mix, following manufacturer’s protocol. The PCR amplified DNA products were confirmed by sequencing (GENEWIZ^®^,https://www.genewiz.com).

### Lentivirus construction and cell transduction

2^nd^ generation lentiviral transfer constructs containing the SFFV promoter upstream of CFTR with and without the transgene separated by a self-cleaving T2A peptide packaging plasmid (pMD2.G) and envelop plasmid (pCMVR8.74) were packaged into HEK293T cells via co-transfection with lipofectamine 2000 (Thermofisher) (25). Target cells were incubated with viral particles in suspension with OptiMEM and 8μg/ml polybrene for 6 hours. Viral titre was measured by quantifying GFP positive cells with flow cytometry on an Attune NxT 2 and analysed with FlowJo software.

### Halide-sensitive yellow fluorescent protein (YFP) Quenching Assay

HEK293T cells were transfected with 100ng pcDNA3.1_YFP (H148Q/I152L) and CFTR cDNAs (WT, K978C, hCAI and h^K978C) (Fig. 2A) using lipofectamine (Thermofisher). At 24 hours post transfection, the medium was replaced with 100 μl Standard buffer (*S*I, Table 6).

YFP fluorescence was analysed using ImageXpress, with a 20x objective, and 472 excitation / 520 emission filters. Images were acquired every 2 seconds for 150 seconds. At 20 seconds 100mM Γ was added to the extracellular medium and at 40 seconds, 10 μM forskolin was added. Images were analysed using ImageJ (http://rsbweb.nih.gov/ij/). Anion binding to YFP(H148Q/I152L) decreases fluorescence (F), thus F/Fmax is used to quantify I-entry. CFTR activity was quantified as the maximal rate of I^-^ entry into cells (mM/s) (26, 27).

### Analysis of transgene expression

Cells were analysed for transgene expression 96 hours after transduction. Images of GFP fluorescence were taken on a Cytation™ 5 - Cell Imaging Multi-Mode Reader at 4x magnification. Images were analysed using ImageJ to quantify the fluorescent:non-fluorescent cell percentage. Transduction efficiency was calculated using the following equation, where n is the number of images taken per well:

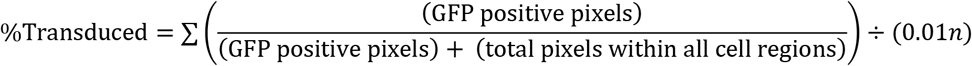

### Western Blots

Cells were lysed in NP-40 Lysis Buffer (*S*I, Table 6), incubated on ice for 20 minutes then centrifuged at 15,000g for 20 minutes. Protein concentration was determined by Pierce™ BCA Protein Assay Kit and 20μg protein was denatured with 2.5μl LDS sample buffer and 1μl reducing agent (NuPAGE) a at 37°C for 30 minutes. Samples were resolved on NuPAGE 4-12% Bis-Tris Protein Gels with mass standards 10-250 kDa (LI-COR). Proteins were transferred to Immobilon^®^-FL PVDF membrane (Millipore). Membranes were blocked in Odyssey^®^ Blocking Buffer (LI-COR), immunostained with anti-CFTR or anti-α Tubulin followed by IRDye^®^ 800CW Goat anti-Mouse IgG (LI-COR), visualised and quantified on an Odyssey IR imager (LI-COR) (*S*I, Tables 4 and 5)

### Fluorescence recovery after photobleaching (FRAP)

GFP linked CFTR transduced CFBE BMI-1 were imaged on a Leica SP5 inverted confocal microscope 63X/1.3 glycerol as described (28). Regions of interest (ROIs) of 20 pixels were selected as the point of highest fluorescence of individual cells. The mean fluorescence of 3 scan iterations (~1 second per iteration) were acquired. ROIs were using 100% transmission of the 488-nm-wavelength laser and fluorescence recovery after photobleaching (FRAP) was measured for ~70 scan iterations. The fluorescence values were normalised to the initial pre-bleach value (1) and the value immediately post-bleach (0). The Mobile Fraction (Mf) was calculated using the equation: 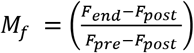

Pre-FRAP images were used to obtain mean fluorescence values from the intracellular region of the cell. At least 10 regions of interest (ROI) were measured for each cell. Ratio of intracellular to peripheral fluorescence were obtained using NIS-Elements software.

### Statistics

Normally distributed data were analysed using ANOVA followed by Tukey’s test or unpaired t-test with Welch’s correction. Paired t tests were applied to samples from the same donor but subject to different treatments. Non-parametric equivalents (Mann–Whitney test, Kruskal–Wallis test with Dunn’s multiple comparisons test) were used when data were not normally distributed. Data are shown as individual points and/or mean ± standard deviation. For linear regression, the coefficient of determination (R^2^) and whether the slope significantly differed from zero are presented. ns: no significant difference. Significant differences are indicated with *: p<0.05; **: p<0.01; ***: p<0.001. Data analyses were performed using GraphPad Prism v9.1.0 (GraphPad Software).

## Acknowledgments

Funded by the Cystic Fibrosis Trust Project No: SRC 006. Provision of cells and media was supported by TARRAN17GO from the Cystic Fibrosis Foundation, BOUCHE15RO from the Cystic Fibrosis Foundation and P30 DK065988 from the NIH, USA.

We thank the CF SRC team for advice and input. Particularly, Ileana Guerrini (ICH) for help in developing the lentiviral constructs. We thank the University of North Carolina (UNC) Cystic Fibrosis Center Tissue Core (director: Scott Randell) for providing cells, media and expert advice. In addition, we thank Rhianna Lee, Michael Chua, Lolita Radet, Saira Ahmad, Patrick Moore, Megan Webster, Ozge Beyazcicek, Eric Scott and Maria Sassano for their help and support at UNC.

