## Supplementary Information for "K978C CFTR restores essential epithelial function with greater efficiency than wildtype CFTR when expressed in CF airway cells"

**This PDF file includes:**

Figures S1 to S4
Tables S1 to S6

**Supporting Information**

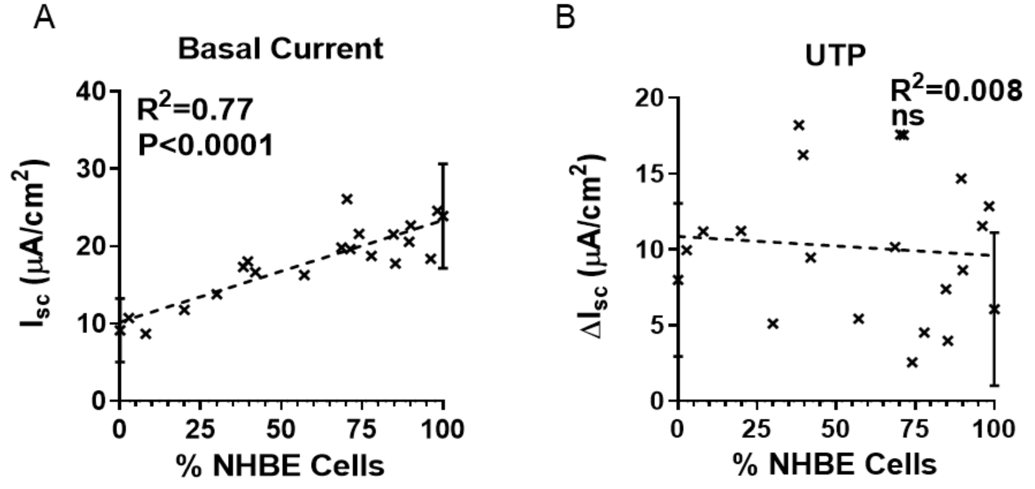


**Fig. S1.** $I_{sc}$ plotted against % NHBE:CFBE (n=8-12 from 3 CF donors for (*A*) basal or (*B*) $\Delta I_{sc}$ after addition of UTP (10 μM). UTP was added apically. Data were subject to linear regression. The $R^2$ values and P value for slope significantly different from zero, are shown on the graphs. ns., not significantly different.

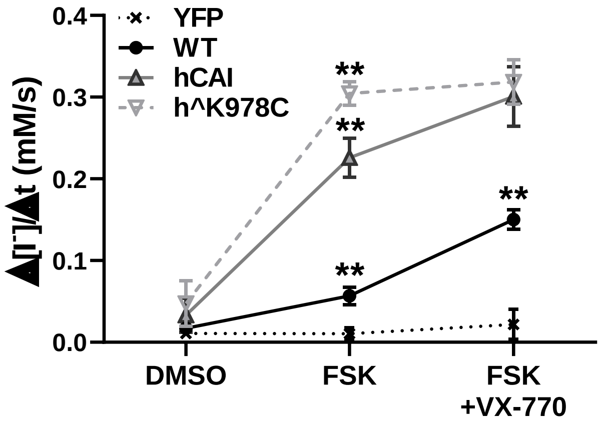


**Fig. S2.** The maximal rate of $I^-$ entry ($\Delta[I^-]/\Delta t$) summarised for CFTR cDNA and conditions. Data are shown as mean values points and means +/- SD. Treatments were compared by two-way ANOVA with Tukey's post hoc analyses; Significantly different as shown *: p<0.05; **: p<0.01; *** :p<0.001, n=3

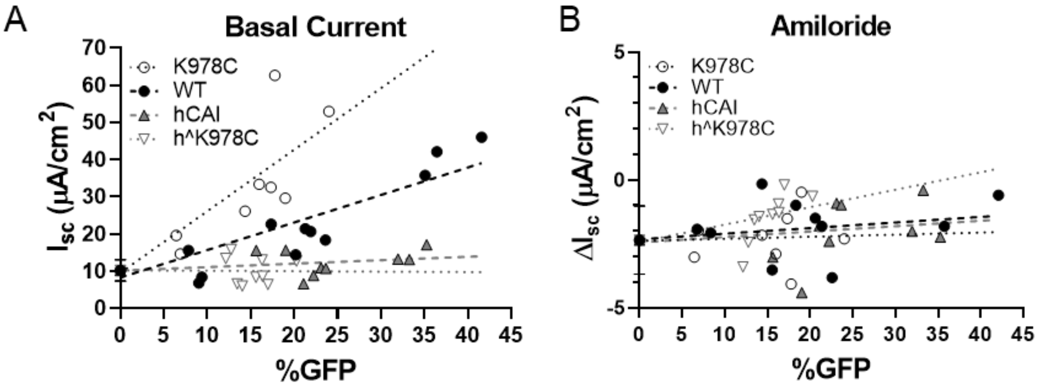


| | K978C | WT | hCAI | h^K978C |
|---|---|---|---|---|
| R square | 0.79 | 0.82 | 0.15 | 0.00 |
| Slope | 1.66* | 0.74* | 0.09 | 0.00* |

| | K978C | WT | hCAI | h^K978C |
|---|---|---|---|---|
| R square | 0.00 | 0.07 | 0.05 | 0.20 |
| Slope | 0.01* | 0.02 | 0.02* | 0.07* |

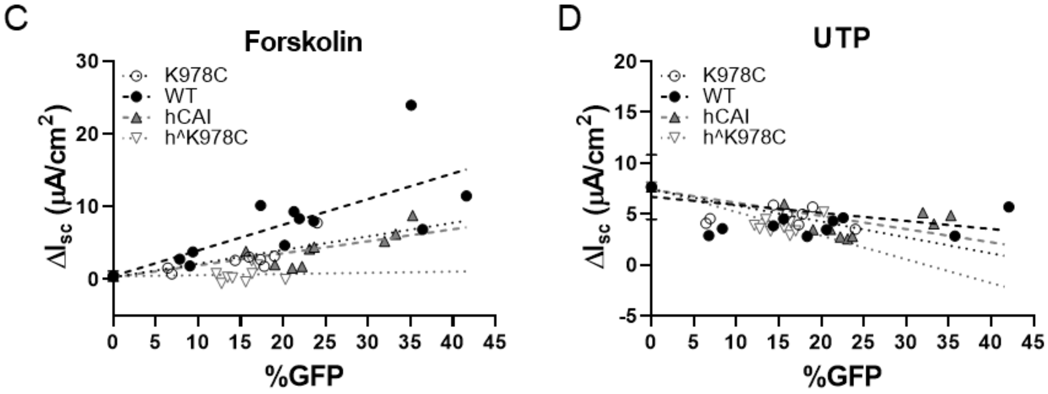


| | K978C | WT | hCAI | h^K978C |
|---|---|---|---|---|
| R square | 0.70 | 0.70 | 0.79 | 0.02 |
| Slope | 0.19* | 0.35* | 0.17* | 0.02* |

| | K978C | WT | hCAI | h^K978C |
|---|---|---|---|---|
| R square | 0.24 | 0.16 | 0.33 | 0.39 |
| Slope | -0.16* | -0.08 | -0.13* | -0.23* |

**Fig. S3.** $I_{sc}$ plotted against plotted against % transduced for each CFTR cDNA for (*A*) basal or (*B*) $\Delta I_{sc}$ after addition of amiloride (10 µM), (*C*) forskolin (10 µM) and (*D*) UTP (10 µM). All drugs were added apically except forskolin which was bilateral. Dotted lines show linear regression with $R^2$ values and significance values for variation of the slope from zero for each CFTR cDNA are shown in the table below *: p<0.05; **: p<0.01; ***: p<0.001. (n=8-12 from 3 CF donors).

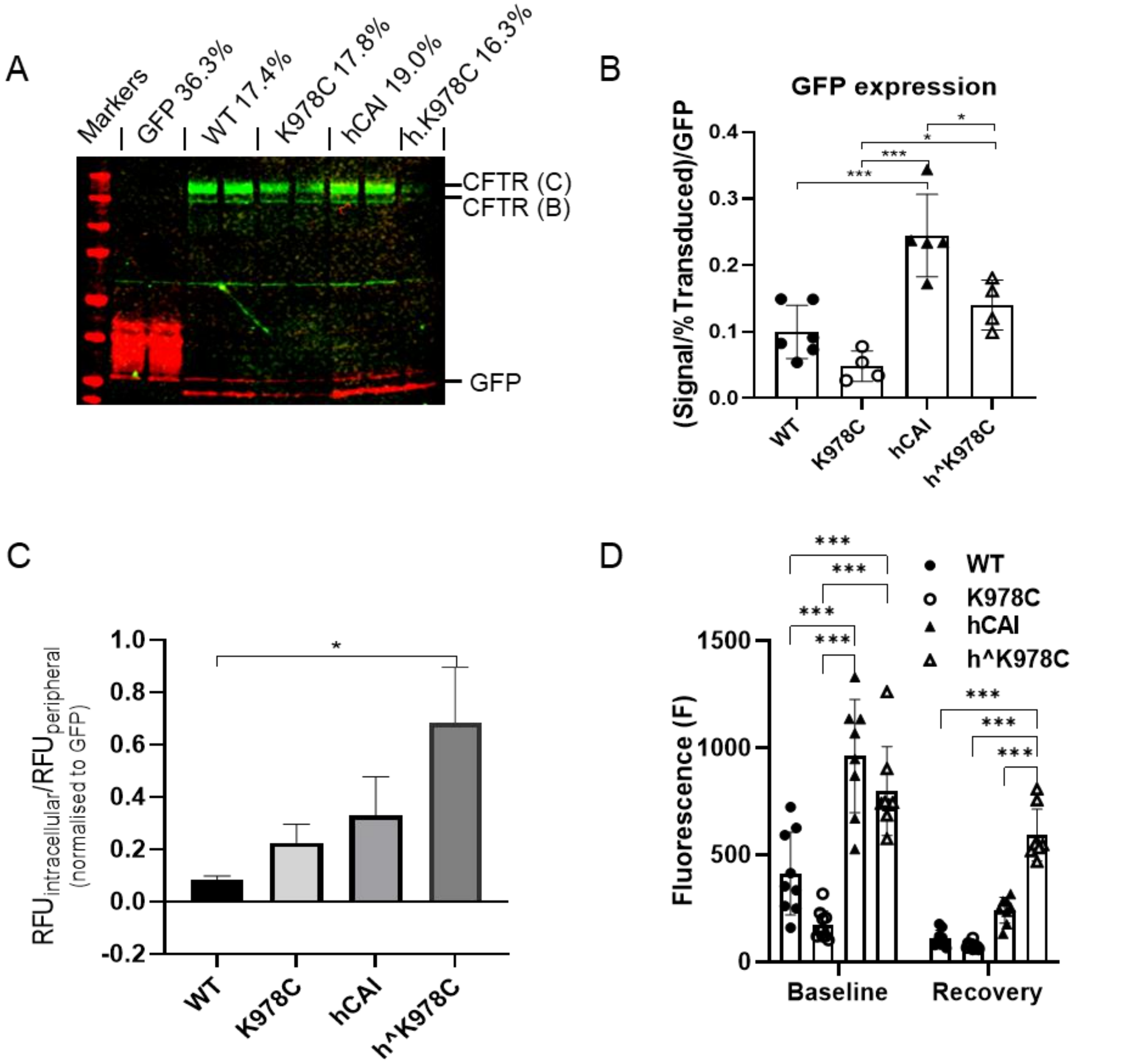


**Fig. S4.** (A) Full size western blot (as in figure 4) of lysate soluble fraction from CFBE cultures transduced with different CFTR cDNAs or GFP alone (as labelled above wells and run in duplicate). Blots were immunostained with anti-CFTR (CFTR 596) and anti-GFP. CFTR protein bands C and B (green), GFP (red) with molecular weight markers shown to the left of the blot (red). (B). Summary of analysis of relative fluorescence units (RFU) for GFP bands relative to % transduced and normalised to GFP only transduced for each CFTR cDNA. Presented as mean ± SD (n=4-7 from 3 donors). Treatments were compared by one-way ANOVA with Tukey's post hoc analyses; Significantly different * $p<0.05$, *** $p<0.001$. (C) Quantification of the ratio of cytosolic GFP RFU to peripheral GFP RFU for each transduction, normalised to cytosolic GFP in cells transduced with GFP. Data are shown as mean +/- SD (n=2-5 from 1 CF donor). CFTR cDNAs were compared to WT CFTR by one-way ANOVA with Kruskal-Wallis post hoc analyses. Significantly different as shown; *: $p<0.05$. (D) Summary of fluorescence (F) from region of interest (ROI) of greatest fluorescence on individual cells pre-bleach (baseline) and 70 seconds after bleach (recovery).

**Table S1.** Demographics of human bronchial epithelial cell donors (from figure 1).

| | Donor | Age at collection | Sex | Ethnicity | Genotype | Smoking history |
|---|---|---|---|---|---|---|
| Normal (Non-CF) | 1 | 49 | M | Caucasian | N/A | <1PY |
| | 2 | 30 | M | Caucasian | N/A | non-smoker |
| | 3 | 43 | M | Caucasian | N/A | non-smoker |
| | 4 | 25 | F | Caucasian | N/A | non-smoker |
| CF | 1 | 26 | M | Caucasian | delF508/delF508 | non-smoker |
| | 2 | 28 | F | Caucasian | delF508/delF508 | non-smoker |
| | 3 | 28 | F | Unknown | delF508/delF508 | non-smoker |
| | 4 | 36 | F | Caucasian | delF508/delF508 | non-smoker |
| | 5 | 28 | M | Unknown | delF508/delF508 | non-smoker |
| | 6 | 34 | F | Unknown | W1282X/R1162X | non-smoker |
| | 7 | 41 | M | Unknown | W1282X/R1162X | non-smoker |

<IPY, less than one pack per year smoking history.

**Table S2.** Demographics of induced sputum donors (pooled for CF sputum).

| | Donor | Age at sputum collection | Sex | Ethnicity | $FEV_1$ | FVC | Genotype |
|---|---|---|---|---|---|---|---|
| CF sputum samples | **1** | 35 | F | Caucasian | 1.21 | 2.28 | DF508/DF508 |
| | **2** | 19 | F | Caucasian | 2.89 | 4.02 | DF508/2184insA |
| | **3** | 59 | M | Caucasian | 0.92 | 2.46 | 3849+10kbC>T/3659delC |
| | **4** | 27 | M | Caucasian | 1.29 | 2.40 | DF508/DF508 |
| | **5** | 29 | M | Caucasian | 2.38 | 4.46 | DF508/DF508 |
| | **6** | 36 | F | Caucasian | 1.21 | 2.35 | DF508/DF508 |
| | **7** | 33 | M | Caucasian | 2.27 | 3.30 | DF058/G451V |
| | **8** | 24 | M | Caucasian | 1.32 | 3.33 | DF058/S945L |
| | **9** | 38 | M | Caucasian | 1.00 | 2.00 | DF508/DF508 |
| | **10** | 59 | M | Caucasian | 1.09 | 2.38 | 3849+10kbC>T/3659delC |

**Table S3.** Primary antisera.

| Name | Species | Dilution | Catalogue # | Supplier |
|---|---|---|---|---|
| Anti-a-Tubulin, clone YL1/2 | Rat | 3 µg/mL | mab1864 | Sigma-Aldrich |
| Anti-CFTR 596 | Mouse | 1:2000 | A4 | Cystic Fibrosis Foundation Therapeutics |
| Anti-GFP - ChIP Grade | Rabbit | 1:5000 | ab290 | Abcam |

**Table S4.** Secondary antisera and fluorophores.

| Name | Species | Fluorophore | Dilution | Catalogue # | Supplier |
|---|---|---|---|---|---|
| Anti-Mouse IgG | Donkey | Alexa Fluor 594 | 1:1000 | A32744 | Thermo Fisher Scientific |
| Anti-Goat IgG | Donkey | Alexa Fluor 555 | 1:1000 | A32816 | Thermo Fisher Scientific |
| Anti-Rabbit IgG | Donkey | Alexa Fluor 488 | 1:1000 | 711-545-152 | Jackson ImmunoResearch |
| Phalloidin | - | Alexa Fluor 647 | 1:50 | A22287 | Thermo Fisher Scientific |
| DAPI | - | Ex/Em 340/ 488 nm | 1:200 | D9542 | Sigma-Aldrich |
| IRDye® anti-Mouse IgG | Goat | 800CW | 1:20000 | 926-32232 | LI-COR Biosciences |

**Table S5.** Buffer composition.

| Solution | Composition |
|---|---|
| NP-40 lysis buffer | 25 mM Tris-HCL pH 7.4, 150mM NaCl, 1mM EDTA, 1% NP-40, 5% Glycerol. Add 1 tablet/10ml cOmplete™, Protease Inhibitor Cocktail at use |
| Ussing buffer | 117 mM NaCl, 2.5 mM $CaCl_2$, 4.7 mM KCl, 1.2 mM $MgSO_4$, 25 mM $NaHCO_3$, 1.2 mM $KH_2PO_4$, 11 mM D-glucose, 5 mM Hepes (pH 7.4) |
| $Cl^-$ free Ussing buffer | 115 mM Na-isethionate, 25 mM $NaHCO_3$, 3mM Ca-gluconate, 2.4mM Mg-gluconate, 2.4mM $K_2HPO_4$, 1.1 mM $KH_2PO$, 11 mM D-glucose, 5 mM Hepes (pH 7.4) |
| Blocking buffer | 1% BSA, 1% fish gelatin, 0.1% Triton X-100, 5% normal goat serum (this can be replaced with BSA) in 1x TBS. |
| Standard buffer | 140 mM NaCl, 4.7 mM KCl, 1.2 mM MgCl2, 5 mM HEPES, 2.5 mM CaCl2, 11 mM glucose, pH 7.4. |
| DNA lysis buffer | 10%SDS, 50 mM EDTA, 50 mM Tris-HCL, 100 mM NaCl, 5 mM DTT, 0.5 mM spermidine. Add Proteinase K 1:100 at use. |

**Table S6.** Primers.

| Name | Sequence 5'-3' | Supplier |
|---|---|---|
| K987C_F | GTGGGATTCTTAATAGATTCTCCTGCGATATAGCA ATTTTGGATGACCTTC | Sigma-Aldrich |
| K978C_R | GAAGGTCATCCAAAATTGCTATATCGCAGGAGAA TCTATTAAGAATCCCAC | Sigma-Aldrich |
| h^K987C_F | GCAGGAGGAATACTAAATAGATTTAGTTGCGATAT AGCAATACTAGATGATCTACTACC | Sigma-Aldrich |
| h^K978C_R | GGTAGTAGATCATCTAGTATTGCTATATCGCAACT AAATCTATTTAGTATTCCTCCTGC | Sigma-Aldrich |
| T2A_P16A_F | CGTGGAGGAGAATGCGGGCCCTATGCAGC | Sigma-Aldrich |
| T2A_P16A_R | GCTGCATAGGGCCCGCATTCTCCTCCACG | Sigma-Aldrich |
| AMEL_F | CCCTGGGCTCTGTAAAGAATAGTG | Amplification |
| AMEL_R | CAGGCTTGAGGCCAACCAT | Amplification |
| Amelogenin-X isoform (AMELX+FAM) | 6-FAM-ATCCCAGATGTTTCTCAA-MGB-NFQ Designed by (George *et al.*, 2013) | ddPCR |
| Amelogenin-Y isoform (AMELY+VIC) | VIC-CATCCCAAATAAAGTGGTT-MGN-NFQ Designed by (George *et al.*, 2013) | ddPCR |
